# Genetic architecture of behavioural resilience to ocean acidification

**DOI:** 10.1101/2022.10.18.512656

**Authors:** Robert Lehmann, Celia Schunter, Megan J. Welch, Stefan T. Arold, Göran E. Nilsson, Jesper N. Tegner, Philip L. Munday, Timothy Ravasi

## Abstract

Genetic variation is essential for adaptation to rapid environmental changes. Identifying genetic variation associated with climate-change related phenotypes is therefore the necessary first step towards predictive models of genomic vulnerability.

Here we used a whole-genome scan to identify candidate genetic variants associated with differences in behavioural resilience to ocean acidification in a coral reef fish. We identified three genomic regions that differ between individuals that are behaviourally tolerant compared with behaviourally sensitive to elevated CO_2_. These include a dopamine receptor (*drd4rs*), cadherin related family member 5-like (*cdhr5l*), Synapse-associated protein 1 (*syap1*), and GRB2 Associated Regulator of MAPK1 Subtype 2 (*garem2*), which have previously been found to modify behaviour related to boldness, novelty seeking, and learning in other species, and differ between behaviourally tolerant and sensitive individuals.

Consequently, the identified genes are promising candidates in the search of the genetic underpinnings and adaptive potential of behavioural resilience to ocean acidification in fishes.

## Introduction

Anthropogenic stressors are impacting the physiology, ecology, and behaviour of marine and terrestrial animals at a global scale (Poloczanska et al. 2013; Buxton et al. 2017; Hendry et al. 2008). Yet the response of individual organisms or populations to different environments is not uniform, with significant variation in traits ranging from the camouflage of walking stick insects (Farkas et al. 2013), foraging behaviour of salamanders (Urban 2013) to the body composition of copepods (Charette and Derry 2016). Instead, there is intraspecific phenotypic variation conferring higher fitness of some individuals in altered environmental conditions. In the dark-eyed Junco, for example, populations inhabiting regions with higher thermal heterogeneity also show increased flexibility in thermogenic capacity compared to populations in thermally homogenous regions (Stager et al. 2021). Indeed, the impact of such intraspecific phenotypic trait variation on a range of ecological response variables is expected to be similar to phenotypic variation across species (Des Roches et al. 2017). If intraspecific phenotypic variation has a large genetic component it can be the basis for adaptation (Falconer and Mackay 1996) to climate change (Bitter et al. 2019; Yang et al. 2021) or other anthropogenic stressors (Biro and Post 2008; Arlinghaus et al. 2017) via natural selection. Therefore, identifying the genetic variation associated with differences in individual fitness in different environments is crucial to making accurate predictions about the biological impacts of climate change and other anthropogenic stressors over the timeframes at which they are occurring (Hoffmann and Sgrò 2011; Razgour et al. 2019; Munday et al. 2013b). In the yellow warbler, genetic variation close to the genes *DRD4* and *DEAF1*, linked to exploratory and novelty-seeking behaviour in several species, was found to be important for successful climate adaptation and allowed the construction of a predictive model for this species (Bay et al. 2018). Modern genomics approaches are making it possible to directly identify candidate allelic variants of genes or loci that can be the raw material for genetic adaptation (Waldvogel et al. 2020) and, therefore, might enable species to adapt to rapid environmental change.

Ocean acidification, caused by the uptake of additional carbon dioxide from the atmosphere, has diverse effects on marine species, including decreased survivorship, altered metabolism, reduced calcification, growth and development (Wittmann and Pörtner 2013; Kroeker et al. 2013; Kelly and Hofmann 2013). In addition, elevated carbon dioxide partial pressure (pCO_2_) has been linked to behavioural changes in some fish and invertebrates, with broad-ranging effects on sensory systems, learning and decision making (Paula et al. 2019; Cattano et al. 2018; Wang and Wang 2020; Munday et al. 2019; Heuer and Grosell 2014). These behavioural changes have been linked to impaired function of GABA_A_ neurotransmitter receptors in the brain (Nilsson et al. 2012; Thomas et al. 2020; Schunter et al. 2019; Heuer et al. 2016; Chivers et al. 2014), as a consequence of acid-base regulation to defend tissue pH against the acidifying effects of high pCO_2_.

One behavioural effect observed in fish exposed to elevated CO_2_ is an altered response to olfactory cues, including an impaired response to the chemical cues of predators and chemical alarm cues (CAC) from conspecifics (Munday et al. 2010; Nilsson et al. 2012; Porteus et al. 2018; Williams et al. 2019; Ferrari et al. 2011; Ou et al. 2015). Alarm cues are chemicals released from the skin of injured prey, which reliably signal high predation risk to other individuals of the same species (Chivers and Smith 1998). Failure to respond appropriately to predator odor or CAC increases the risk of predation (Ferrari et al. 2010). Indeed, field-based experiments show that larval reef fishes exposed to elevated CO_2_ for 4-5 days, which induced impaired responses to predator odor and CAC, exhibit markedly higher rates of mortality from predation in their natural habitat (Munday et al. 2010; Ferrari et al. 2011; Chivers et al. 2014). Nevertheless, variation in behavioural sensitivity to elevated CO_2_ has been detected, with some individuals displaying greater tolerance to elevated CO_2_ than others (Welch and Munday 2017; Schunter et al. 2016; Munday et al. 2013a). While the behavioural response of some individuals to predator odor or CAC is impaired at CO_2_ levels predicted to occur by the end of the century (700-800 μatm), the behaviour of other individuals is unaffected (Munday et al. 2010, 2013b; Ferrari et al. 2011; Welch et al. 2014). In the spiny damselfish, *Acanthochromis polyacanthus*, variation in behavioural response to CAC under elevated pCO_2_ is correlated in fathers and their offspring, suggesting a heritable genetic basis (Welch and Munday 2017), which might enable populations of this species to adapt to rising CO_2_ levels in the ocean. The transcriptional brain response of juvenile *A. polyacanthus* to high CO_2_ has been studied, showing differential expression patterns between the offspring of behaviourally tolerant or sensitive parents (Schunter et al. 2016, 2018), but the genetic variants associated with the observed variation in behavioral tolerance to elevated CO_2_ remain unknown.

Here, we used a whole-genome scan to compare the genotype of *A. polyacanthus* individuals that have a strongly impaired behavioural response to conspecific alarm cue (i.e. are behaviourally sensitive) compared with the genotype of individuals that show a typical response (i.e. are behaviourally tolerant) to CAC under elevated CO_2_ conditions. To do this we re-sequenced the genomes of fish reared in conjunction with the study of Welch and Munday (Welch and Munday 2017) to test the heritability of behavioural tolerance to elevated CO_2_ in *A. polyacanthus*. Briefly, wild-caught adult fish from the same population were exposed to elevated CO_2_ (754 μatm, consistent with climate change projections) and their behavioural response to CAC was tested, assigning individuals retaining a strong natural avoidance of CAC to the tolerant group and individuals being attracted to CAC to the sensitive group. Breeding pairs were formed from similar-sized males and females in the behaviourally tolerant group and from similar-sized males and females in the behaviourally sensitive group. Offspring from these tolerant and sensitive breeding pairs were then reared under control or elevated CO_2_ conditions for up to five months. In the current study, the genomes of adult fish (N=18) together with a large number of their offspring (N=210) were then re-sequenced to generate a high confidence set of genetic variants. Comparing this genetic variant set in the adult fish that had been categorized as either tolerant or sensitive to elevated CO_2_ allowed us to identify genomic candidate regions and genes that differentiate individuals that are behaviourally tolerant or sensitive to elevated CO_2_ (Fig. 1a).

**Figure 1:**
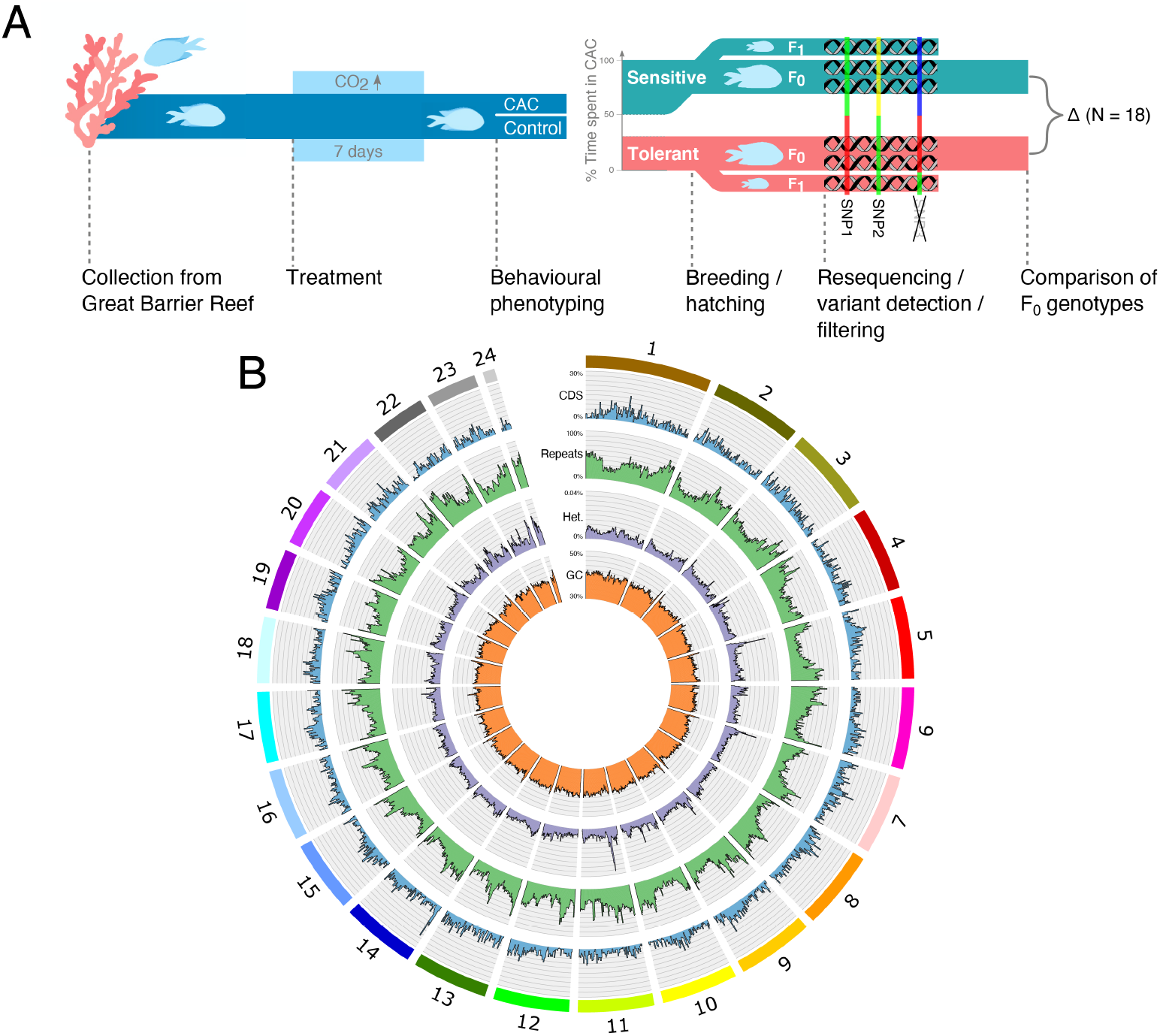
Experiment setup and the reference genome assembly. **a)** Experimental design with the F_1_ rearing treatments elaborated in Fig. S1 **b)** The genome assembly of *A. polyacanthus*, consisting of 24 pseudo-chromosomes (outermost ring), with coding sequence density (blue), repetitive sequence density (green), heterozygosity in non-repetitive sequence (purple), and GC content (orange) shown for non-overlapping windows of 500 kb width.

## Results & Discussion

### A chromosome-scale reference genome for *Acanthochromis polyacanthus*

A whole-genome scan for genetic differentiation requires a high-quality reference genome. We generated a long-read dataset with 131x coverage of brain genome from an adult *Acanthochromis polyacanthus* using the SMRT sequencing platform of PacBio and assembled a high quality highly contiguous brain reference genome assembly, which was then placed in chromosome-scale scaffolds using a chromatin contact map from Hi-C data (see Tables S1, S2, S3, and Figures S2, S3). The resulting *A. polyacanthus* genome assembly consists of 25,468 annotated genes on 24 pseudo-chromosomes (Fig. 1b). It features an N50 of 41.7 Mb with 96 % of the initial assembly being placed in chromosomes and 57 unplaced contigs. The assembly includes more transposable element insertion sites, leading to an increased repeat content of 38 % (Fig. S4, Table S4), compared to a previous short-read based assembly (Schunter et al. 2016) as well as a higher assembly level completeness (BUSCO score 96.7 %) with reduced duplication (Table S5). Assembly of the mitochondrial genome and construction of a phylogeny using mitochondrial and nuclear marker genes also confirms the species of the sequenced individual (Fig. S5, Table S6).

### Genomic regions segregating with behavioural phenotype

To identify a set of polymorphic single nucleotide polymorphisms (SNPs) in the sample population of *A. polyacanthus* we re-sequenced the genomes of 18 behaviourally tested adult individuals. In addition, we re-sequenced 210 offspring of these adults to allow for pedigree-based filtering later on. Sequencing yielded an averaged 103.5 million raw reads per individual resulting in 32.7x coverage (Table S7). Comparison of all re-sequenced individuals to the reference genome assembly allowed for the identification of 16.4 million polymorphic genomic sites. The pedigree information available for the 210 offspring allowed for rigorous quality filtering resulting in a final set of 5.88 million SNPs (Table S8). This high-quality set of variants was then used to identify genomic loci linked to the behavioural phenotype in the eighteen adults that had been behaviourally tested. For this we used the cross-population haplotype-based statistic XP-nSL (Szpiech et al. 2020), cross-population extended haplotype homozygosity XP-EHH (Sabeti et al. 2007), and the fixation index F_ST_ in a windowed genome-wide scan (Fig. 2, see Methods for details on window definition and filtering). Similar analyses with comparable number of samples have been used previously to identify genomic regions associated with tame behaviour in farm-bred red foxes (Kukekova et al. 2018), genomic signatures of speciation for in Lake Victoria cichlids (Nakamura et al. 2021) and North American songbirds (Termignoni-Garcia et al. 2022) amongst others. This analysis yielded three genomic outlier regions with consistently strong signal for all three measures (Table S9, Fig. 2). Region I located in the center of chromosome one contains the strongest differentiation signal found in coding sequences across the whole genome, with large positive values for XP-nSL and XP-EHH, specifically in the protocadherin gene *cdhr5l* and the dopamine receptor D4 (*drd4rs*) (Fig. 2 bottom). The positive value of XP-nSL and XP-EHH indicates longer homozygous haplotypes in the tolerant cohort, suggesting a shared origin. In contrast, Region II on chromosome five features significantly negative XP-nSL and XP-EHH values (Fig. S10), which suggests a shared origin of the sensitive haplotype that is centered around the gene coding for the Synapse-associated protein 1 (*syap1*). Region III on chromosome 16 also features negative XP-nSL and XP-EHH values and is centered around GRB2 associated regulator of MAPK1 subtype 2 (*garem2*) (Fig. S11). In summary, the behaviourally tolerant cohort features a determining haplotype at the *drd4rs*/*cdhr5l* locus while the behaviourally sensitive cohort is characterized by two determining haplotypes at the *syap1* and *garem2* loci.

**Figure 2:**
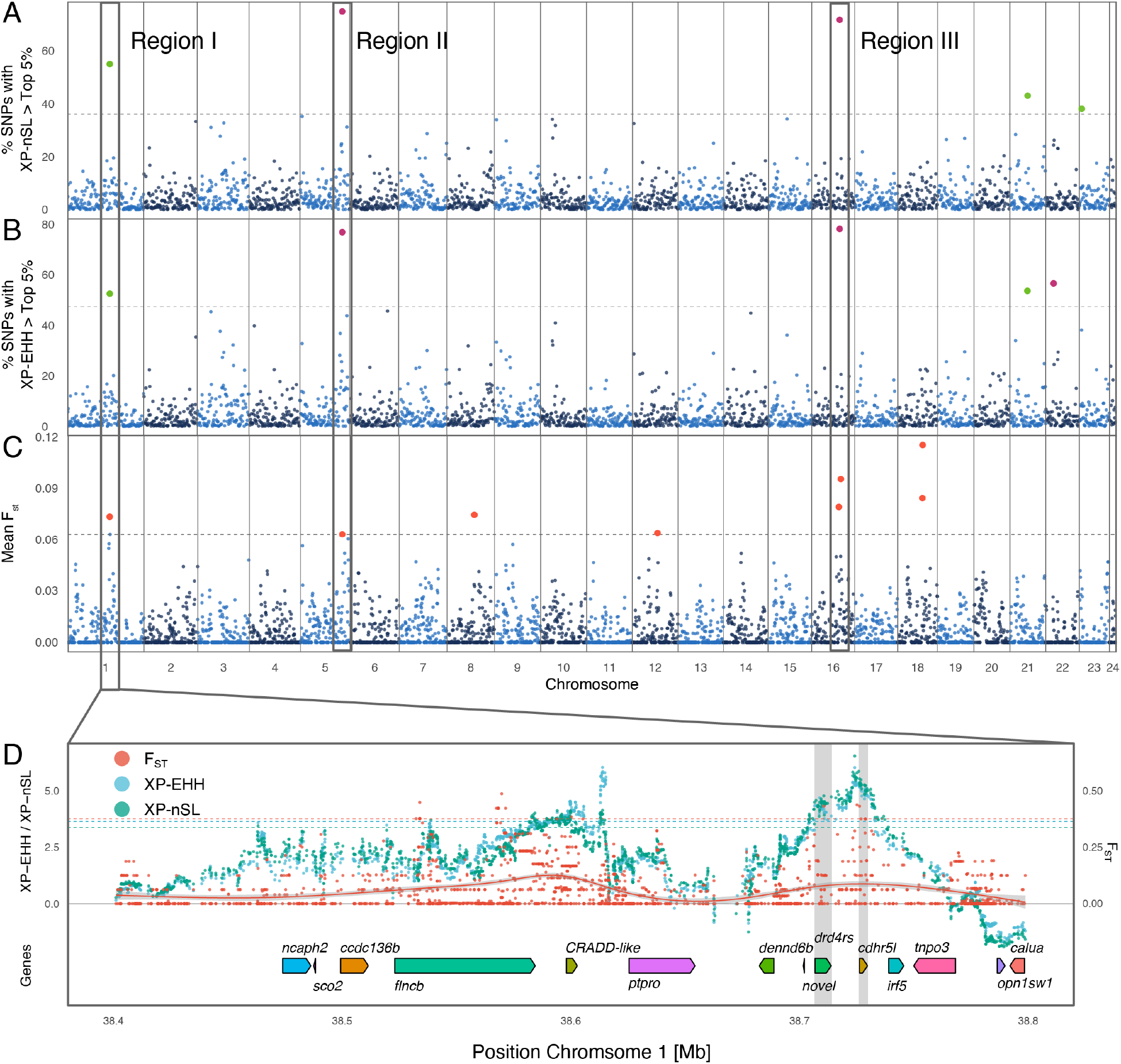
Genome-wide scan for genetic differentiation between behaviourally tolerant and sensitive fish to elevated CO_2_. Density of SNPs with extreme a) cross-population nSL (XP-nSL) and b) cross-population extended haplotype homozygosity score (XP-EHH). Each point represents a genomic window of 400 kB width for which the percentage of SNPs among the top 5% most extreme values genome-wide is shown. c) Mean fixation index F_ST_ within 500 kB genomic windows. The dotted lines mark the top 0.2 % cutoff for each measure across all windows, outlier windows are marked in green for positive XP measures, purple for negative XP measures, and orange for F_ST_. Three regions marked I to III are consistent outliers across measures. d) Detailed comparison of SNP-wise differentiation measures in genomic region I together with annotated genes and the genomic loci of *drd4rs* and *cdhr5l* are shaded in gray. A local polynomial fit of F_ST_ values is shown as a solid line.

### Dopamine receptor 4 (*drd4rs*) shows signature of genetic differentiation

Region I (Fig. 2) contains the two genes with the strongest differentiation signal (Table S9) including one of three dopamine D4 receptors (*drd4rs*, mean XP-EHH 4.21, mean XP-nSL 4.38) and its neighboring gene cadherin-related family member 5-like (*cdhr5l*, mean XP-EHH 4.95, mean XP-nSL 5.13). The local polynomial fit of the fixation index F_ST_ follows the pattern of the haplotype-based measures with two peaks of positive value, suggesting the causal haplotype in the tolerant cohort. The G-protein coupled receptor gene *drd4rs* of *A. polyacanthus* (1,238 bp) consists of 4 exons that code for 7 transmembrane domains and one long intracellular loop (Fig. 3a). A SNP in the third exon, encoding the fifth transmembrane domain (TM5) segregates with the behavioural phenotype (F_ST_ 0.43, top 0.0004 percentile) while inducing a synonymous codon change.

**Figure 3:**
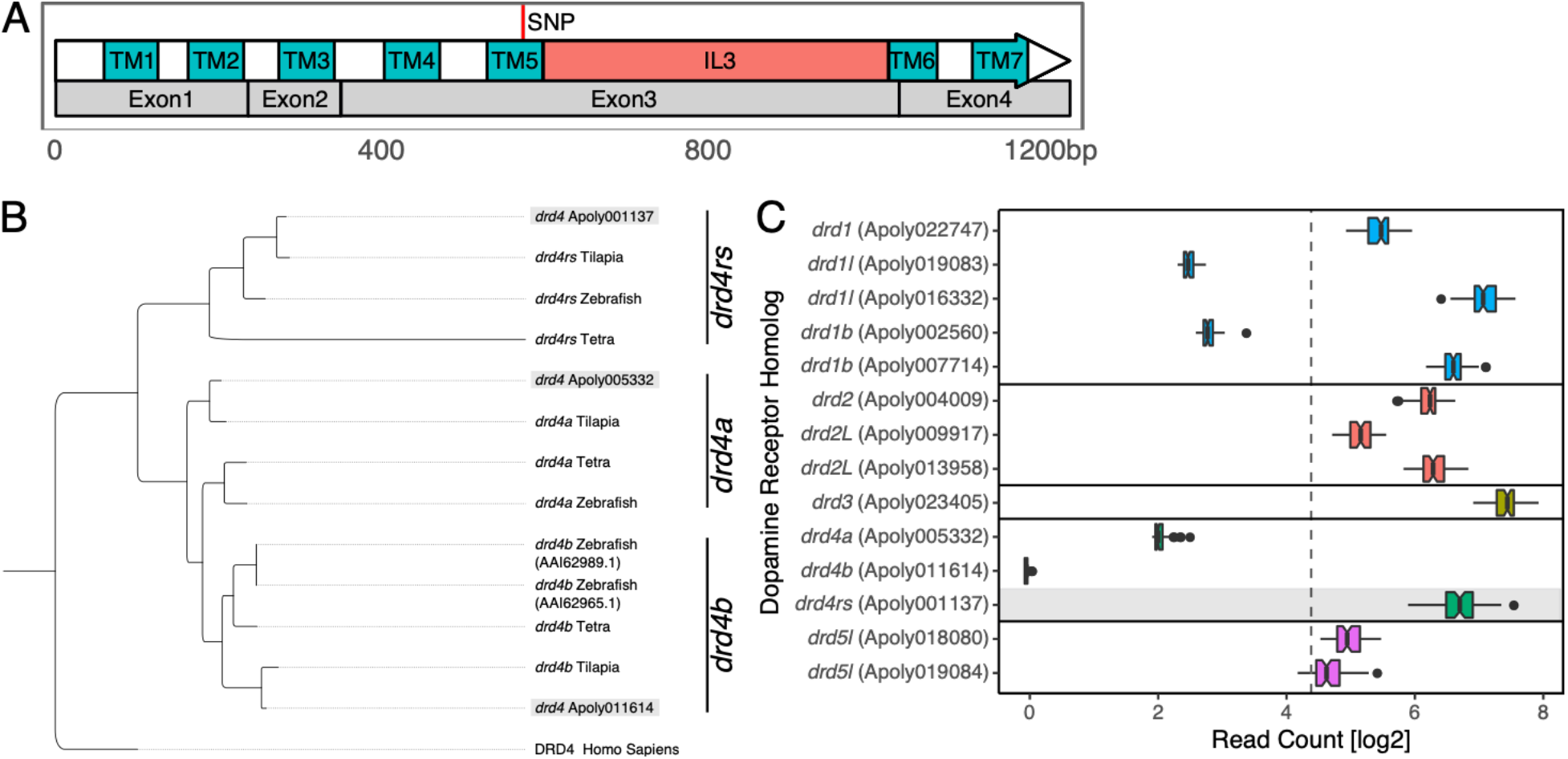
Domain structure of Dopamine Receptor D4 (*drd4rs*) and phylogenetic and transcriptional analysis of dopamine receptor genes. **a)** A SNP segregating with the behavioural phenotype is located in the third exon of *drd4rs*, encoding the fifth transmembrane domain (TM5) prior to the intracellular loop 3 (IL3). **b)**Phylogenetic tree of *drd4* homologs from our study species, *A. polyacanthus*, and *Oreochromis niloticus*, *Astyanax mexicanus*, and *Danio rerio*, using the human D4 receptor gene as outgroup. **c)**Gene expression levels of all members of the dopamine receptor family (mean normalized read count including all samples from all CO_2_ treatments). The mean dataset-wide expression is marked as gray line while gene boxes are color-coded by receptor subgroup 1 to 5.

In birds, synonymous SNPs in *drd4* are associated with behavioural traits such as novelty-seeking or escape behaviour from a cage (Mueller et al. 2014; Kluen et al. 2012; Mueller et al. 2013). Specifically, two synonymous SNPs in *drd4* exon 3 of invasive populations of the Yellow-crowned bishop explain 11% and 15% of the neophobic and neophilic behavioural phenotypes for two populations, respectively (Mueller et al. 2014). In blue tit populations, a synonymous exon 3 SNP was associated with escape behaviour from a cage (Kluen et al. 2012). In some great tit populations one synonymous SNP in *drd4* exon 3 was found to be associated with behaviour in a novel environment chamber (Mueller et al. 2013). Furthermore, genetic variation upstream of the *drd4* gene of the yellow warbler appears to be highly relevant to its ecology. This locus was found to feature one of the highest associations between genotype and climate across a range of populations, for which the genotype information was then used to predict genomic vulnerability to climate change (Bay et al. 2018). Taken together, these results demonstrate that in wild bird populations, genetic polymorphisms in the dopamine 4 receptor gene are at least partially responsible for setting the baseline behavioural response to avoidance inducing stimuli, suggesting that the observed synonymous variant in the *drd4rs* gene of *A. polyacanthus* might similarly modify the typically avoidant response to alarm cues when exposed to elevated CO_2_.

The family of dopamine receptor genes is separated into two major groups (group 1: *drd1* and *5* and group 2: *drd2*, *3*, and *4*) (Opazo et al. 2018). While mammals and birds generally only have one *drd4* gene, teleosts feature three homologous D4 receptor genes a, b, and rs. Through phylogenetic analysis we ascertained that it is the drd4rs which is carrying the identified genetic variation (Fig. 3b). To evaluate if there is active transcription of this or other candidate genes in the brain of *A. polyacanthus*, we re-analyzed previously published transcriptomics data (Schunter et al. 2018) of 72 offspring exposed to elevated CO_2_ conditions for various lengths of time from behaviourally tolerant and sensitive breeding parents (Table S12). Briefly, these CO_2_ conditions included transgenerational exposure of parents and offspring to elevated CO_2_, developmental exposure of offspring to elevated CO_2_ from hatching, and acute exposure to elevated CO_2_ for four days in five months old offspring (see Fig. S1 and Methods for more details). Comparison of dopamine receptor family members shows *drd4a* and *b* to be extremely low or not expressed while *drd4rs* is expressed constitutively across all individuals and treatments (Fig. 3c), which is consistent with the possible activity of *drd4rs* in the brain.

### D4 receptors might modulate aversive behavioural response signaling via indirect pathway in D2 receptors

In order to respond to alarm cue stimulus different environmental inputs (Leahy et al. 2011) are integrated before inducing the avoidance behaviour. This points towards the involvement of the basal ganglia, a part of the forebrain that plays a central role in motivational and cognitive learning, where the so-called direct and indirect pathways mediate the response to positive and negative stimuli (Albin et al. 1989). An aversion stimulus, such as a chemical alarm cue, decreases dopaminergic signaling which reduces the induction of direct pathway neurons, the D1 receptors, and de-repression of indirect pathway neurons, which are the dopamine receptors of the D2 family to which D4 receptors belong (Albin et al. 1989). It was demonstrated experimentally that suppressing firing of dopaminergic neurons in the ventral tegmental area of mice is sufficient to activate the indirect pathway via D2 receptors, inducing aversive behaviour and learning (Danjo et al. 2014). Dopamine receptors of type D4 were shown to form heterodimers with D2 receptors, thereby modulating their output in the striatum in humans (Borroto-Escuela et al. 2011; González et al. 2012). Specifically, dopaminergic signaling is increased upon reduced D2-D4 heteromerization, in this case due to a structural variation in the D4 receptor gene (Bonaventura et al. 2017; Sánchez-Soto et al. 2018). The comparison of the protein structure of the *A. polyacanthus drd4rs* gene to the human D4 receptor protein reveals a well conserved dopamine binding pocket structure (Fig. 4a, S13) and a long unstructured third intracellular loop (IL3) mediating the heteromerization of the D4 receptor with other receptors (Woods 2010), supporting functional homology between both. While the observed genetic differences between behaviourally tolerant and sensitive individuals do not reflect an amino acid change in the D4 protein in *A. polyacanthus*, this difference might still have functional consequences. Synonymous SNPs in the D2 gene were found to modify the expression and stability of the mRNA and thus influence the protein abundance (Duan et al. 2003) by changing the ribosomal pausing propensity (McCarthy et al. 2017), explaining the observed association of seemingly silent SNPs to a psychiatric disorder in humans. Although gene expression based on whole-brain tissue of this gene did not differ in our experiment on *A. polyacanthus*, it is possible that the observed non-coding SNP in the *drd4rs* gene differentiating tolerant and sensitive individuals might alter the protein levels in certain brain regions. This could lead to an alteration of D2-D4 receptor heterodimers and a modified D2-dependent indirect pathway activation. Behavioural phenotypes fitting this hypothesis were found in experimental pharmacological perturbations of dopaminergic signaling in *Danio rerio*. Amongst the observed phenotypes were swimming distance (Ek et al. 2016), occurrence of swimming episodes (Thirumalai and Cline 2008), activity (Tran et al. 2015), dark-phase activity (Shontz et al. 2018), and notably hypoactivity in response to a high affinity D4 receptor antagonist (Boehmler et al. 2007). Similarly, an increased expression of the D2 receptor resulted in bolder behaviour in *Danio rerio* when placed in a novel tank (Thörnqvist et al. 2019). The involvement of the direct/indirect pathway circuitry in fish is supported by the extensive homology of brain regions between *Danio rerio* and mammals (Parker et al. 2013). Furthermore, a study of cleaner wrasse (*Labroides dimidiatus*) and client fish (*Naso elegans*) interactions under elevated CO_2_ conditions found a large decrease in dopamine in the midbrain of the cleaner wrasse, which was associated with decreased cleaning interactions (Paula et al. 2019). Hence, there is evidence from a range of model systems, including fish, that alterations in dopaminergic signaling can lead to similar boldness-related behavioural phenotypes as observed in *A. polyacanthus*. Furthermore, synonymous SNPs can also lead to alterations in dopamine receptor abundance, potentially causing an imbalance between dopamine receptor homologs and thereby disturbing dopaminergic signaling.

**Figure 4:**
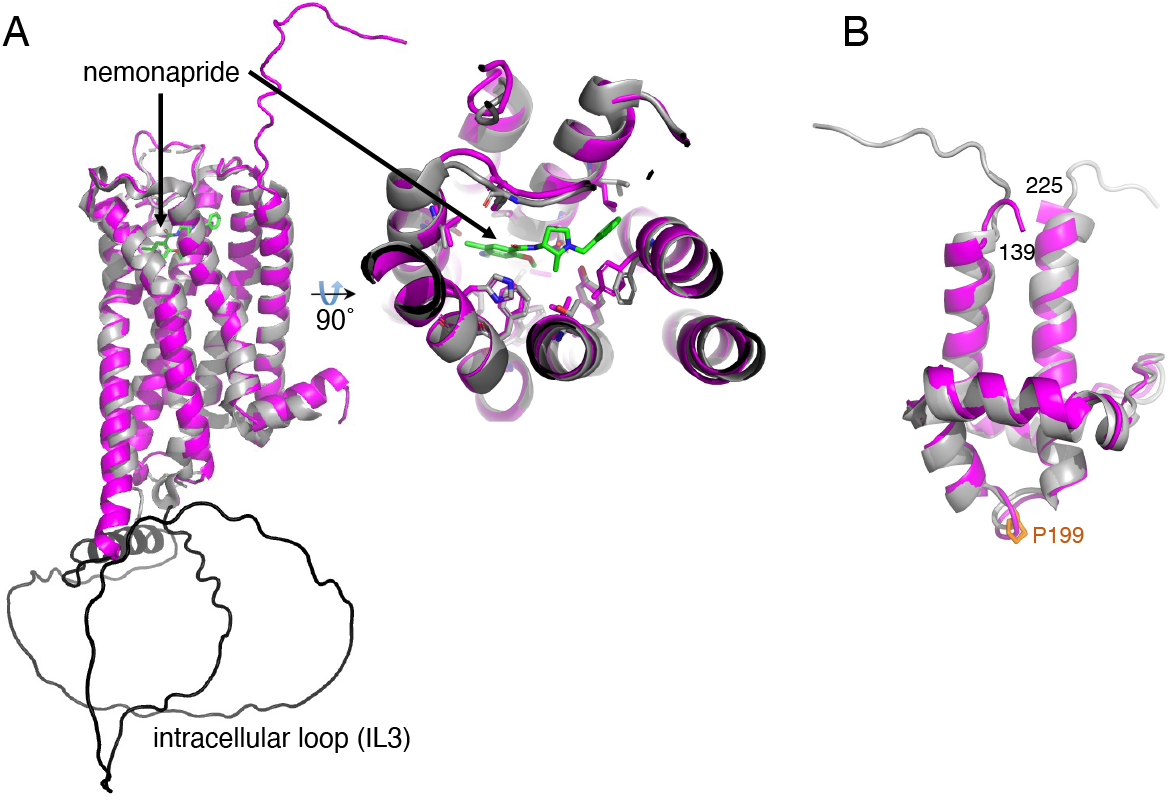
Protein structure prediction of candidate genes segregating behavioural phenotypes. **a)** The AlphaFold predicted structure of the *A. polyacanthus drd4rs* gene (pink) superimposes with an RMSD of 0.8 Å to the human D4 receptor gene (grey, PDB Id: 5wiu). The selective antagonist nemonapride is shown as a green stick model. The long predicted intracellular loop IL3 is shown in black. On the right, the view into the dopamine binding pocket reveals ligand-binding residues, which are nearly all strictly conserved (key residues and nemonapride are shown as stick models). **b)** Structural AlphaFold model of the BSD domain of the *syap1* Leu199 variant (magenta; only residues 139-225 are shown), superimposed onto the human Synapse-associated protein 1 BSD domain structure (grey; RMSD is 0.7 Å; PDB accession 1×3a) featuring a highly conserved proline in this position (shown as orange stick model).

Prior research has demonstrated the involvement of GABA_A_ receptors in the behavioural response to olfactory stimuli under elevated CO_2_ in coral reef and other marine fishes (Nilsson et al. 2012; Chivers et al. 2014; Williams et al. 2019; Schunter et al. 2019). The synaptic homeostasis of both D1 and D2 receptor-expressing neurons is maintained by extra-synaptic GABA_A_ receptors in mice, which modulate their excitability by induction of a tonic current in response to prolonged activation (Maguire et al. 2014). Moreover, the activity of the ventral tegmental area has been found to be modulated via dense GABAergic projections (Barrot et al. 2012) in response to stimuli and three distinct fear and anxiety-related behaviours were reduced upon introduction of lesions in this area (Jhou et al. 2009). This interweaving of dopaminergic and GABAergic signaling is consistent with the observation that perturbation of GABAergic signaling with gabazine can reverse the high CO_2_-induced behavioural phenotype (Nilsson et al. 2012; Chivers et al. 2014). It is thus tempting to suggest that the genetic variation we see in the dopamine D4 receptor *drd4rs* of *A. polyacanthus* modifies the behavioural response to a stimulus by altering the sensitivity balance between the direct and indirect pathway. This variation gives rise to bolder individuals unlikely to respond aversely to the alarm cue stimulus under elevated CO_2_ and less bold individuals who are more likely to respond aversely to alarm cue.

### Possible association of fish-specific protocadherin to behavioural variation

Next to the dopamine receptor, the gene with the strongest differentiation signal between tolerant and sensitive behavioural phenotypes under elevated CO_2_ in *A. polyacanthus* is cadherin related family member 5-like (*cdhr5l*, Apoly001138) (Fig. 2). It is part of the superfamily of cadherin genes which is comprised of several large gene families (classical cadherins, desmosomal cadherins and protocadherins) with diverse functions in cell-cell interaction during development, differentiation, migration, axon outgrowth, dendrite arborization, synapse formation, stabilization, and plasticity (Seong et al. 2015). *CDHR5* is classified as non-clustered protocadherin group epsilon according to Kim *et al.* (Kim et al. 2011) due to the four cadherin repeat domains (human *CDHR5* as well as *A. polyacanthus*’ *cdhr5l* Apoly001138). We found a significantly differentiated synonymous SNP featuring the highest fixation index (F_ST_ 0.43, top 0.0004 percentile), while one non-synonymous SNP (F_ST_ 0.31, top 0.0024 percentile) and two successive SNPs close to the splice acceptor site of exon 10 also feature significant differentiation (F_ST_ 0.375, top 0.001 percentile). The non-synonymous SNP Met348Thr is located within the third cadherin repeat domain that is part of the interface to cadherin repeat 4 (Fig. S9), where it codes either for a methionine or threonine amino acid. Cadherins are transmembrane proteins which mediate cell-cell adhesion via their extracellular cadherin domains. These cadherin domains need to be stabilised in an extended position by calcium ions that are lodging between successive cadherin domains to stabilise their linker regions. The substitution of the long hydrophobic methionine with a shorter and partly hydrophilic threonine is expected to affect the associations of this position with the core of the cadherin repeat 3, and thus the position of the loop containing arginines 89 and 90. Such a conformational change in one of the extracellular cadherin repeat domains may influence the responsiveness and dynamics of the cadherin to calcium, and hence affect its capability and dynamics of cell-cell interaction mediation between neurons in the brain.

The differentiating *cdhr5l* Apoly001138 as well as its homolog Apoly011615 are highly expressed in the *A. polyacanthus* transgenerational elevated CO_2_ treatment (Fig. S12). Furthermore, various members of this gene family are expressed in the brain and differentially regulated in response to different CO_2_ treatments. In mammalian model organisms non-clustered *PCDH* are predominantly expressed in the nervous system and are important for the establishment of selective synaptic connections between the cerebral cortex and other brain regions, such as the thalamus (Kim et al. 2007), and the maintenance and plasticity of the adult hippocampus (Kim et al. 2010). Similarly, protocadherins exhibit complex expression patterns in the brain of *Danio rerio*, where they are required for normal functioning and maintenance (Liu et al. 2015). A recent study showed that cadherin mediates the stabilization and long term potentiation of excitatory synapses at dopaminergic neurons in the ventral tegmental area (Mills et al. 2017). Perhaps unsurprisingly then, mutations in various protocadherins have been linked to a wide range of behavioural disorders in humans (Tsai and Huber 2017). These results suggest a potential role of this still unstudied protocadherin homolog *cdhr5l* in the maintenance and plasticity of brain regions upstream of the basal ganglia of *A. polyacanthus*, allowing for the possibility that the observed genetic variation might impact the behavioural response to alarm cue under elevated CO_2_.

### Synapse associated protein 1 of region II is associated with synaptic plasticity-related behavioural phenotypes in various model systems

We found three paralogs of Synapse-associated protein 1 (*SYAP1)*, expressed at low levels in the brain of offspring *A. polyacanthus* (Fig. S10), with one harboring a non-synonymous SNP at amino acid position 199 (either a proline or a leucine) in the functionally relevant BSD domain, which segregates with the behaviourally tolerant and sensitive phenotypes (F_st_ 0.28, top 0.005 percentile). Similar to the tolerant cohort, the experimental structure of the 80% identical BSD domain from human Syap1 features a proline in *cis* conformation in position 199 (PDB accession 1×3a). This *cis*-bonded proline allows the α4-α4 loop to connect both helices while sealing the small hydrophobic core (Fig. 4b). Prolines are energetically more favorable and hence >100 times more likely to form *cis* bonds than all other residues (Joseph et al. 2012). Therefore, substituting Pro199 with a leucine as found in the sensitive cohort is expected to alter the α4-α4 loop conformation and to destabilize the BSD domain. The function of BSD domains is unknown, but prolines are highly conserved at this position (Doerks et al. 2002), suggesting that this *cis*-proline is functionally relevant. *Cis*-*trans* changes are associated with the evolution of new functions (Joseph et al. 2012), supporting a role of this variant in fish behavior.

Support for a possible role of *syap1* in the response to alarm cue is found in fruit fly and mouse model systems. Firstly, larvae of Sap47 knockout fruit flies exhibit deficiencies in short-term plasticity involved in olfactory associative memory processing (Saumweber et al. 2011). Specifically, these larvae show a ~50% reduction in the ability to learn and/or remember the association of an odorant with a rewarding tastant. Furthermore, recent work demonstrated that Sap47 knockout reduces the lifespan, impairs climbing proficiency, and reduces the plasticity in circadian rhythm and sleep (Blanco-Redondo et al. 2019). Secondly, in mice *SYAP1* is prominently expressed in the nervous system where a knockout reduces locomotor activity in early phases when voluntary movement is initiated (Von Collenberg et al. 2019). Accordingly, the observed genetic variation in *A. polyacanthus* might modulate the protein functional efficiency and thereby olfactory learning and/or voluntary movement initiation in response to alarm cue in elevated CO_2_.

### *garem2* in region III is a modifier of anxiety-like behavior in mice

The third region significantly differentiating between the two cohorts contains a member of the Grb2-associated and regulator of Erk/MAPK gene family (*GAREM*) (Tashiro et al. 2009). *A. polyacanthus* features one copy of *garem2*, which is expressed in the brain of offspring from both phenotypes, with a mean-variance stabilized expression level of 8.2 across all samples. Furthermore, we observe significant expression induction under developmentally elevated CO_2_ treatment of the offspring (log2-fold-change 0.34, p < 2.08E-6). Two SNPs in the first intron exhibit a high F_ST_ of 0.38 (top 0.0009 percentile) as well as a set of SNPs upstream (Fig. S11), while six variants within the coding sequence do not segregate with the phenotype (F_ST_ < 0.0625, below top 0.12 percentile). Prior work in mice has shown the involvement of *GAREM2* with behaviour similar to the alarm cue response. Knocking out *GAREM2* in mice leads to a reduction of anxiety-like behavior, increased social approaching and exploratory behavior, and a reduction in novelty-induced anxiety (Nishino et al. 2019).

### Candidate loci for genetic predisposition of behavioural tolerance to elevated CO_2_ stressor

Currently, the genetic structure responsible for the phenotypic variation in behavioral resilience to ocean acidification is not known. Here we present a whole-genome scan to detect genomic candidate loci that might be responsible for this variation. We find variation in the genes coding for dopamine receptor D4 (*drd4rs*), cadherin related family member 5-like (*cdhr5l*), Synapse-associated protein 1 (*syap1*), and GRB-associated and regulator of ERK/Mapk subtype 2 (*garem2*) that correlate with behaviourally tolerant and sensitive phenotypes expressed under elevated CO_2_ conditions. Each of these genes has been connected to modifications of boldness, exploratory, and anxiety-related behavioural patterns in a range of model species, supporting their role in the impairment of behavioural response to risk cues (e.g. alarm cues and predator cues) in *A. polyacanthus* and other fishes (Welch et al. 2014; Nagelkerken and Munday 2016; Cattano et al. 2018; Williams et al. 2019; Munday et al. 2019; Porteus et al. 2018). The family of dopamine receptors is mechanistically well studied, leading to the hypothesis that the observed genetic variation in behaviourally tolerant *A. polyacanthus* modifies the aversive behavior signaling in response to negative stimuli in the basal ganglia, thereby expanding the acclimatization range compared to sensitive individuals. The genetic variation in *cdhr5l*, *syap1* and *garem2* of sensitive individuals might have the opposite effect upstream of the basal ganglia, collectively modifying the baseline behavioural response of an individual to a negative stimulus, such as conspecific alarm cues. Elevated environmental CO_2_ modifies the genetically predisposed typical behavior, rendering sensitive individuals less responsive to alarm cue, while tolerant individuals manage to respond appropriately, as they would under normal CO_2_ conditions.

Our results suggest a link between the alarm cue avoidance behaviour to well-described signaling mechanisms during aversive behaviour in several model organisms and furthermore suggest that there is standing genetic variation in key behaviour-associated genes that would provide the raw material for adaptation of behavioural responses in *A. polyacanthus*, and probably other fishes, to rising CO_2_ levels in the ocean. Our results also constitute the ideal starting point for further validation using quantitative genetics, pharmacological perturbation, or genome editing to elucidate the cellular mechanisms responsible for altered behavioural responses to elevated CO_2_ and the genetic variation that could foster genetic adaptation of marine animal behaviour to ocean acidification.

## Methods

### Specimen collection and experimental design

A total of 121 adult spiny damselfish *Acanthochromis polyacanthus* collected in the central Great Barrier Reef (GBR), Australia (18°38’24,3” S, 146°29’31,8” E) were exposed to 754 ± 92 μatm CO_2_, a level projected to occur by the end of this century (Collins et al. 2013), for 7 days as previously described (Welch and Munday 2017). The 7 day exposure duration was chosen because previous studies show that fish display impaired responses to chemical stimuli after four or more days of exposure to elevated CO_2_ (Ferrari et al. 2011; Heuer et al. 2016; Munday et al. 2010). A two-chamber flume was used to determine the behavioural phenotype of these 121 individuals in response to conspecific chemical alarm cue (CAC). A ratio of one donor fish per test fish was used in the preparation of CAC. The CAC was obtained by euthanizing the donor fish with a quick blow to the head, making superficial incisions on both sides of the body, rinsing the body with 60 ml of control water, and adding that to 10 l of seawater with elevated CO_2_. The flume was fed with control and CAC water at a constant rate of 450 ml/min from two header tanks (one with 10 l control water and one with 10 l CAC treated water). A fresh preparation of CAC treated water was used for each test fish. Individuals were subjected to nine minute long behavioural trials consisting of two minutes habituation, a two minute recording period, one minute switching of CAC and control sides, followed by another two minutes habituation and two minutes recording period. The position of the fish in the flume was noted every five seconds during the two x two minute recording periods. Large variations in the behavioural response to CAC were observed, with some individuals avoiding the CAC as they normally do in ambient conditions, whereas other individuals preferred CAC over control water. The former were termed behaviourally ‘tolerant’ (< 30 % of the time in CAC) while the latter were termed behaviourally ‘sensitive’ (> 50 % of time in CAC) to elevated CO_2_. Since the experiment was designed to answer a range of complementary questions pertaining to the heritability of the observed behavioural phenotype, including epigenetic effects, it included generating a generation of offspring from the phenotyped adult individuals. The offspring from this experiment that were collected for molecular analyses were not behaviourally phenotyped, and are thus not informative for the comparison of tolerant vs. sensitive individuals, but their genotypes are useful to filter the SNP panel which is why they were included here. Breeding pairs were then assigned based on sex and size into four different groups: (1) tolerant male and tolerant female, (2) sensitive male and sensitive female, (3) tolerant male and sensitive female, and (4) sensitive male and tolerant female. Only the first two groupings, comprising both tolerant or both sensitive individuals are considered in this study. The latter two “mixed phenotype” groups were used in (Welch and Munday 2017), but are not included in the current study. Half of the tolerant and sensitive breeding pairs in each group were allowed to acclimate to control condition and half to elevated CO_2_ conditions for three months prior to the start of the breeding season. A total of five tolerant and five sensitive pairs bred in control conditions, and a total of four tolerant and five sensitive pairs bred in elevated CO_2_ conditions. To reduce any potential family related bias, two parental pairs from each behavioural phenotype were allowed to breed in control condition and then moved to elevated CO_2_, acclimated for three months and bred in elevated CO_2_ (included in counts above). Following breeding, five randomly selected tolerant pairs and five randomly selected sensitive pairs were sacrificed for extraction of brain tissue (Table S1). This sums to a total of 20 parental fish, but the total number of sequenced adults was 18 due to DNA degradation observed in four samples and the addition of two available phenotyped samples not selected for breeding pairs. The average percentage time in CAC of the sampled/sequenced tolerant adults was 3.6 % (SD 10.3) and the average percentage time in CAC of the sampled/sequenced tolerant adults was 87 % (SD 18.2).

On hatching, offspring clutches from the breeding pairs were immediately moved to separate tanks maintaining the CO_2_ conditions of the respective parental pair, where they were reared for up to five months. In addition, some offspring reared under control conditions were exposed to elevated CO_2_ levels for four days prior to sampling. These combinations resulted in four different treatments: 1. (Control) parents and offspring under control CO_2_ level, 2. (Transgenerational) parents and offspring under elevated CO_2_ level, 3. (Developmental) parents under control CO_2_ level while offspring are exposed to elevated CO_2_ immediately after hatching, and 4. (Acute) parents and offspring are exposed to control CO_2_ levels, but with a four-day long exposure of offspring to elevated CO_2_ levels before sampling (Fig. S1). Offspring environmental condition exposures were maintained for two durations, five weeks and five months, after which individuals were euthanized. This yielded 228 individuals (18 adults and 210 offspring) suitable for DNA sequencing: F_0_ adult nine tolerant / nine sensitive, F_1_ five weeks old: 102, F_1_ five months old: 108. From this experiment brains of 72 five months old offspring were previously extracted for RNA and sequenced (see Schunter et al. 2018) and re-analyzed in this study. The experiment was conducted under James Cook University ethics approval A1828.

### DNA extraction, library preparation, and sequencing for *de novo* genome assembly

To build a reference genome sequence, we collected one large adult fish from the wild at Bramble reef on the Great Barrier Reef, Australia. This individual was not phenotyped since the reference does not influence the result of a genome scan contrasting the sensitive and the tolerant group, as neither are required to match the reference. The whole brain tissue was dissected, snap-frozen in liquid nitrogen and kept at −80°C prior to processing. High molecular weight DNA was then extracted from this tissue using the Qiagen Genomic-tip 100/G extraction kit. Briefly, after homogenization of the whole brain tissue using sterile beads and lysis buffer G2 supplemented with 200 μg/mL RNase A for 30 sec, proteinase K was added followed by overnight incubation at 50°C. DNA was then extracted following the protocol resulting in a final elution volume of 200 μl. Pulsed-field gel electrophoresis was used to assess DNA fragment size and quality. The extracted DNA was sheared with a g-TUBE (Covaris, MA, USA) to a target size of 20 kb prior to the preparation of SMRTbell libraries according to the protocol provided by Pacific Biosciences, CA, USA. BluePippin pulse-field gel electrophoresis (Sage Science, MA, USA) was used to perform a fragment size selection to obtain one library with a minimum size of 10 kb and one with 5kb, which were then sequenced using a PacBio RS II instrument at the King Abdullah University of Science and Technology (KAUST) Bioscience Core Laboratory using P6-C4 chemistry and 109 SMRT cells.

From our controlled aquarium experiment, the whole brain tissue was dissected from offspring, snap-frozen with liquid nitrogen and kept at −80°C prior to processing. For the adult individuals, fin clips were taken from the dorsal fins and kept in ethanol for further processing. For genome resequencing, the DNA and RNA of the 228 samples were extracted using a Qiagen AllPrep DNA/RNA Mini Kit. For DNA extraction from the 18 adult finclips a Qiagen Blood and Tissue kit was used. Approximately 10 RNase and DNase free one-use silica beads (Daintree Scientific, Australia) were placed into Eppendorf tubes together with the tissue samples, which were then homogenized for 30 seconds in a pre-frozen metal tray with a Thermo Fisher Scientific bead beater. DNA and total RNA were then purified according to the manufacturers’ protocol and stored at −80 ºC. Illumina sequencing libraries for 49 samples were produced with a TruSeq DNA library preparation kits and sequenced on the HiSeq4000 platform by Macrogen (Macrogen South Korea). An additional 179 samples were sequenced using the same procedure at the King Abdullah University of Science and Technology (KAUST) Bioscience Core Laboratory.

### Genome assembly and proximity guided assembly scaffolding

The PacBio reads (Table S2) were assembled using FALCON v0.4.0 (Chin et al. 2016) varying the parameters generating 11 candidate assemblies (Table S3). These candidate assemblies were compared with respect to contiguity and the best was selected for phasing with FALCON_Unzip and initial polishing with quiver. A chromatin contact map was assembled by Phase Genomics (Seattle, WA, USA). For this procedure, an adult fish was dissected, the flash-frozen brain tissue was fixed and sent to Phase Genomics, where the chromatin was isolated and a library prepared for 80 bp paired-end sequencing. Chromosome-scale scaffolds were then obtained by mapping the read data to the reference assembly, then clustering, ordering, and orienting contigs into 24 clusters (Fig. S2) using Proximo (Bickhart et al. 2017; Burton et al. 2013) as previously described (Peichel et al. 2017). The scaffolded assembly was then polished over three rounds with Arrow, which resulted in the final assembly (Table S3). To confirm the agreement of a short read based genome assembly (GCA_002109545.1) (Schunter et al. 2016) with the new assembly, a whole-genome alignment was performed with Mummer (Kurtz et al. 2004) using default parameters and the result visualized with dotplotly (Fig. S3). This resulted in an alignment of 97.35 % of the short-read assembly to 93.6 % of the new assembly with 99 % average identity.

### Repeat element annotation

A repeat library was constructed with RepeatModeler v1.08 (Smit and Hubley 2008). A second library was constructed with LtrHarvest (Ellinghaus et al. 2008) and LTRdigest (Steinbiss et al. 2009), from the genometools suite 1.5.6 (Gremme et al. 2013) (parameters: -seed 76 -xdrop 7 -mat 2 -mis −2 -ins −3 -del −3-mintsd 4 -maxtsd 20 -minlenltr 100 -maxlenltr 6000 -maxdistltr 25000 -mindistltr 1500 -similar 90). The combined results were deduplicated via clustering with USEARCH (Edgar 2010) (>90% sequence identity) retaining only cluster representatives. The repeat library was then classified by RepeatClassifier. A total of 38% (Table S4) of the genome assembly was masked by RepeatMasker (Smit et al. 2010) using the *de novo* library and the Repbase v22.05 (Bao et al. 2015) *Danio rerio* repeat library. The comparison of transposable element (TE) content between the short-read assembly (annotated using the same *de novo* library, resulting in 25.2 % of sequence masked) and the new long-read assembly reveals that particularly recently inserted TE copies were assembled more successfully (Fig. S4).

### Structural and functional gene annotation

After mapping the RNA-seq data from this previously published experiment (Schunter et al. 2016, 2018) to the final assembly using STAR v2.5.2b (Dobin et al. 2013), BRAKER1 v1.9 (Hoff et al. 2016) was used to perform an *ab-initio* annotation of the soft-masked reference genome assembly, providing the RNA-seq hints and protein sequences of the short read assembly and the closest related fish species with a high-quality reference genome available, the orange clownfish (Lehmann et al. 2019) (GCA_003047355.1) as evidence. This annotation was filtered to obtain a high-quality gene set to train the Augustus (Stanke et al. 2006) gene prediction pipeline. A subsequent BRAKER1 run with similar hints then identified 41,975 genes. The MAKER2 v2.31.8 (Holt and Yandell 2011) gene annotation pipeline was used to annotate the AED score and only genes with an AED < 0.7 were retained for the final annotation (Table S5). InterProScan 5 was executed to obtain the Pfam protein domain. The current UniProtKB/Swiss-Prot, TrEMBL, and NCBI non-redundant database were obtained at 09/2019 and blast 2.6.0 was used to align the annotated protein sequences to these databases, retaining the best hits when falling below an e-value of 1*e^−5^. Annotations in the regions of interest were furthermore validated by mapping protein sequences annotated by Ensembl to the short-read assembly to the new assembly using genomethreader v1.7.0.

### Assembly of mitochondrial genome

The obtained PacBio data were aligned to the mitochondrial genome sequence of the closest related fish species, the orange clownfish *Amphiprion percula* (Lehmann et al. 2019) (CM011763.1). This filtering step retained 466 mapping reads with an N50 of 12,531 bp and 3,997,697 bp total length which corresponds to 240x expected coverage of the assembled mitochondrial genome. This read dataset was then assembled with Organelle_PBA (Soorni et al. 2017) and annotated with MitoAnnotator (Iwasaki et al. 2013) (Fig. S5a). A phylogeny based on the sequences of the ATP synthase 8/6, Cytochrome B (*Cyt b*), and V(D)J recombination-activating protein 1 (*RAG1*) genes from eight damselfish species including another sample of *A. polyacanthus* (Table S6) was then constructed to confirm the species of the sequenced individual. After aligning the concatenated sequences with ClustalW 2.1 (Stamatakis 2006), a maximum-likelihood tree was obtained using RAxML (Larkin et al. 2007) with the GTRGAMMA model and 500 rounds of bootstrapping, affirming the identity of the sequenced individual as *A. polyacanthus* (Fig. S5b).

### Whole-genome resequencing data processing

Illumina short-read sequences obtained from the individuals from the controlled aquarium experiment were assessed with FastQC (Andrews) and low quality regions were trimmed with Trimmomatic v0.33 (Bolger et al. 2014) using parameters: 2:30:10 LEADING:3 TRAILING:3 SLIDINGWINDOW:4:20 MINLEN:40, leaving an average of 97 million reads per sample for analysis (Table S7). Trimmed reads were mapped to the long read assembly and sorted with BWA mem (Li and Durbin 2010), duplicates were removed with sambamba (Tarasov et al. 2015), and read groups added with Picard tools (Broad Institute 2018). Single nucleotide polymorphisms were jointly called in all samples with GATK HaplotypeCaller (McKenna et al. 2010) 3.8.0. An initial high confidence SNP set was generated by filtering the raw variants according to these criteria: QD < 2.0, FS > 60.0, MQ < 40.0, MQRankSum < −12.5, ReadPosRankSum < −8.0, retaining only bi-allelic sites with minimum allele frequency > 0.01, DP > 3, and at least 40 samples with genotype calls. Furthermore, only SNPs conforming to Mendelian inheritance according to our pedigree were retained, yielding 4,296,736 sites (S7 Table, High confidence set). The obtained high confidence variant set was then used as training set for Variant Quality Score Recalibration (VQSR) using these annotations: QD MQRankSum ReadPosRankSum FS MQ SOR DP. The final variant set was then obtained by filtering the VQSR result for an average genotype quality value above 35 and less than 10 violations of Mendelian inheritance per locus. This procedure resulted in a total of 5,882,916 variants (S7 Table, VQSR filtered), which were then phased with shapeit v4.1.3 (Delaneau et al. 2019) using an increased number of iterations with option 10b,1p,1b,1p,1b,1p,1b,1p,10m. Genome-wide distribution of heterozygosity (Fig. 1b) was assessed by identifying repetitive regions with more than three times the expected coverage in each sample, excluding SNPs located in these repetitive regions, and calculating the fraction of remaining heterozygous SNPs in 500 kb non-overlapping windows. PCA (Fig. S6) and Admixture (Alexander et al. 2009) were used to confirm the absence of a population structure signal. The normalized cross-population extended haplotype homozygosity (XP-EHH) and XP-nSL are then calculated between nine sensitive parent individuals and nine tolerant ones using selscan v1.2.0 (Szpiech and Hernandez 2014) setting the sensitive cohort as reference. To define genomic regions that segregate with the behavioural phenotype, the top 5% of SNPs with the most extreme score genome-wide were then defined as outliers as implemented in selscan’s normalization procedure. This resulted in an upper and lower threshold for XP-EHH of 1.95 and −2.04, respectively. Similarly, the upper and lower thresholds for XP-nSL were 1.96 and −2.01, respectively. The genome-wide as well as per-chromosome distributions for SNP-wise XP-EHH and XP-nSL scores are shown in Fig. S7 A-F together with quantile-quantile plots illustrating the defined outlier thresholds. Furthermore, the selscan normalization procedure includes the division of the genome into non-overlapping windows, here set to 400 kb width, where the percentage of outlier SNPs within each window indicates differentiated regions. We selected the top 0.2 % windows with the highest outlier SNP percentage as candidate regions in each differentiation measure, corresponding to 47% and 36% of outlier SNPs per window as thresholds for XP-EHH and XP-nSL, respectively (Fig. S8, Fig. 2 A and B).

The Weir-Cockerham fixation index F_ST_ was calculated between the parental individuals of the same phenotype with vcftools v0.1.16 (Danecek et al. 2011) (see Fig. S7 G for genome-wide distribution), again followed by a windowing into 500kb wide regions with 250kb overlap and calculating the mean F_st_ (Fig. 2C. Again, the top 0.2% of windows with the highest mean F_st_ were selected as candidate regions, corresponding to a threshold of 0.063 (Fig S8 C). Finally, only regions detected as differentiated by all three measures were retained for detailed evaluation to ensure rigorous filtering and avoid false positives. The application of all three genomic differentiation measures was limited to the parental individuals since phenotypes for offspring are not available making their genomic information not informative beyond the SNP filtering step.

The effect of SNPs was estimated with SNPeff (Cingolani et al. 2012).

### Phylogeny construction

A multiple amino acid sequence alignment was constructed using the three *drd4* homologs found in the long-read genome assembly as well as the three *drd4* genes of the Nile tilapia (*Oreochromis niloticus*), Mexican tetra (*Astyanax mexicanus*), and Zebrafish (*Danio rerio*) (Table S11), and lastly the human DRD4 gene (MAFFT using auto setting). A maximum-likelihood phylogenetic tree was obtained from the multiple sequence alignment using RAxML (Larkin et al. 2007) with the model PROTGAMMAAUTO and 500 rounds of bootstrapping.

### RNA-seq data processing

For the RNA-seq read data for 72 samples from the controlled aquarium experiment (Schunter et al. 2016, 2018), data integrity was assessed with FastQC (Andrews), trimmed with Trimmomatic v0.33 (Bolger et al. 2014) (2:30:10 LEADING:3 TRAILING:3 SLIDINGWINDOW:4:20 MINLEN:40) and mapped to the long read assembly with STAR v2.5.2b (Dobin et al. 2013) (Table S12) using the MAKER2 annotation as well as the genomethreader annotation. Differential expression analysis and visualization was then performed with R and the DESeq2 package 1.22.2 (Love et al. 2014). Pairwise differential expression analyses were performed between the acute, developmental, and transgenerational samples against control individuals, separating by parental phenotype.

### Protein structure prediction

Structural models were produced by AlfaFold (Jumper et al. 2021) using the AlphaFold2_advanced.ipynb colab implementation with default values, except for the Filter options (cov: 90; qid: 30). Importantly, this colab version does not use structural templates, and hence is not biased by known PDB structures. Predicted LDDT per-residue scores were above 80 for the secondary structure regions in the transmembrane domain of Drd4 and for the BSD domain of Syap1 (Fig. S13).

## Data Access

All generated data have been deposited at the NCBI under BioProject ID PRJNA671567. Supplementary figures and tables are available at https://doi.org/10.5281/zenodo.7219978.

## Competing Interests

The authors declare no competing interests.

## Acknowledgements

This work was supported by the Office of Competitive Research Funds (OSR-2015-CRG4-2541 and FCC/1/1976-25 to T.R., P.L.M., S.T.A) from the King Abdullah University of Science and Technology, and the Australian Research Council (ARC) as well as the ARC Centre of Excellence for Coral Reef Studies to P.L.M.

## Contributions

M.J.W. and P.L.M. designed and managed the fish rearing experiments. M.J.W. performed the behavioural phenotyping of the adult fish. C.S. prepared the DNA and RNA samples for sequencing, R.L. designed and performed the analyses, S.T.A. performed the protein structure modeling, R.L. wrote the paper with input from C.S., P.L.M., G.E.N., T.R., and J.N.T.

## Notes

https://doi.org/10.5281/zenodo.7219978

### Competing Interest Statement

The authors have declared no competing interest.

